# Cold-Induced Thermogenesis Increases Acetylation on the Brown Fat Proteome and Metabolome

**DOI:** 10.1101/445718

**Authors:** Samuel W. Entwisle, Joan Sanchez-Gurmaches, Robert T. Lawrence, David J. Pedersen, Su Myung Jung, Miguel Martin-Perez, Adilson Guilherme, Michael P. Czech, David A. Guertin, Judit Villen

## Abstract

Stimulating brown adipose tissue (BAT) energy expenditure could be a therapy for obesity and related metabolic diseases. Achieving this requires a systems-level understanding of the biochemical underpinnings of thermogenesis. To identify novel metabolic features of active BAT, we measured protein abundance, protein acetylation, and metabolite levels in BAT isolated from mice living in their thermoneutral zone or in colder environments. We find that the enzymes which synthesize lipids from cytosolic acetyl-coA are among the most robustly increased proteins after cold acclimation, consistent with recent studies highlighting the importance of anabolic *de novo* lipogenesis in BAT. In addition, many mitochondrial proteins are hyperacetylated by cold acclimation, including several sites on UCP1, which may have functional relevance. Metabolomics analysis further reveals cold-dependent increases to acetylated carnitine and several amino acids. This BAT multi-omics resource highlights widespread proteomic and metabolic changes linked to acetyl-CoA synthesis and utilization that may be useful in unraveling the remarkable metabolic properties of active BAT.

## INTRODUCTION

Brown adipose tissue (BAT) specializes in burning calories to generate heat in a process called thermogenesis. Active BAT can rapidly absorb and metabolize glucose and lipids during thermogenesis^1^, and its presence in adult humans correlates with improved metabolic phenotypes^2,3^. Humans with type-2 diabetes (T2D) or obesity have been shown to have lower rates of BAT glucose uptake ^4^, and stimulating BAT activity in mice^5^ and humans^3^, or transplanting BAT into mice^6^, reverses metabolic deficiencies associated with T2D. BAT may also provide metabolic health benefits via secreted factors called BATokines^7-9^. Collectively, these findings suggest that increasing BAT activity may be a promising therapeutic strategy for T2D and obesity in humans. Thus, there is strong rationale for more deeply exploring the biochemical properties of BAT metabolism.

In physiologically relevant settings, BAT is activated by the sympathetic nervous system in response to cold exposure or certain high fat diets^10,11^. Cold-induced thermogenesis (CIT) is the best understood mode of BAT activation. Classically, norepinephrine triggers CIT in brown adipocytes by stimulating β3-adrenergic receptor signaling and cAMP synthesis, which activates protein kinase A (PKA). PKA phosphorylates CREB, which binds to the cAMP response element and induces transcription of many target genes including the uncoupling protein UCP1. UCP1 localizes to the inner mitochondrial membrane and dissipates the proton motive force as heat, thus uncoupling it from ATP synthesis. This is thought to be the primary mechanism of heat-generation in BAT, although alternate heat-generating pathways have been reported^12,13^. While many of the signaling modalities that stimulate BAT have been identified, the associated metabolic and proteomic changes that underlie thermogenesis remain less well defined.

To gain a more comprehensive understanding of the molecular underpinnings of CIT, we analyzed the proteome, acetylproteome, and metabolome of BAT from mice that were acclimated to three different temperatures: thermoneutrality (30°C), room temperature (22°C) and severe cold (6°C). We report striking cold-dependent changes that, when integrated, reveal previously unappreciated changes in acetyl-CoA metabolism and redox control relevant to lipid metabolism and thermogenesis, including acetylation sites on UCP1 that may have functional relevance.

## RESULTS

### Cold adaptation upregulates lipid synthesis, glucose metabolism, and electron transport chain complex I-II proteins

First, we explored proteomic signatures of chronic cold-induced BAT activation by acclimating mice to three different temperature conditions: thermoneutrality (30°C, which is preferred by mice^14^), room temperature (which is mildly cold for mice, 22°C) and severe cold (6°C) (Fig 1A). Following the temperature acclimation period, BAT was harvested and subjected to proteome analysis. The number of confidently detected proteins is consistent across all samples, ranging from 1857 to 2432 (Fig 1B). Differential protein expression was determined by comparing each cold condition to thermoneutrality. We identified 60 proteins altered (∼3%, 26 increasing, 34 decreasing) in response to mild cold and 232 proteins altered (∼11%, 119 increasing, 113 decreasing) in response to severe cold (Fig 1C) indicating a thermogenic proteome response that scales with cold severity.

**Figure 1:**
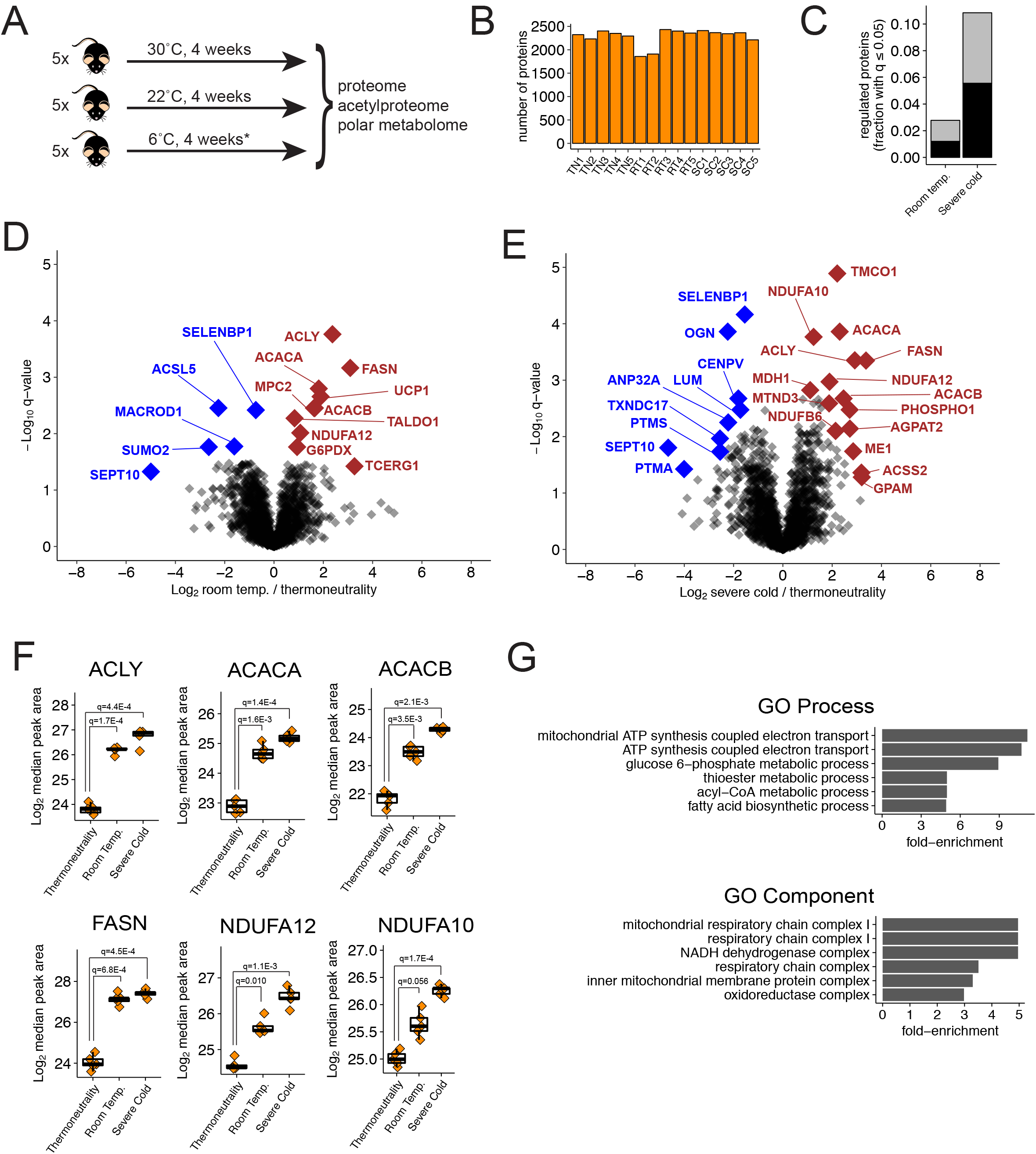
Replicate analysis of BAT after room temperature acclimation reveals a core proteome response of thermogenesis. A. Outline of experimental design. *Temperature was decreased 4°C per week for 4 weeks, such that they were housed at 6°C for the final week. B. Number of identified proteins in each biological condition. Error bars represent standard deviation. C. The fraction of proteins with q ≤ 0.05, either increasing (black) or decreasing (grey), is plotted for the indicated experimental conditions, relative to the thermoneutrality control. D. Volcano plot showing the Log_2_ fold-change versus the -Log_10_ q-value for all proteins, between the severe cold and thermoneutral conditions. Selected increasing (brown) and decreasing proteins (blue) after severe cold acclimation are highlighted. E. Volcano plot showing the Log_2_ fold-change versus the -Log_10_ q-value for all proteins, between the room temperature and thermoneutral conditions. Selected increasing (brown) and decreasing proteins (blue) after severe cold acclimation are highlighted. F. Boxplots showing selected increasing proteins after cold acclimation, in all conditions from both experiments. Data points represent measurements from individual animals. G. Gene Ontology (GO) enrichment analysis of proteins that were significantly increased at least twofold in mild *or* severe cold versus thermoneutrality. The top six most enriched GO Processes and Components are shown, with all enrichments significant at a 5% FDR.

Both mild and severe cold acclimation had profound effects on the proteome. Mild cold acclimation already caused significant increases to *de novo* lipogenesis proteins including ATP citrate lyase (ACLY, 5.2-fold), the two acetyl-CoA carboxylase isoforms (ACACA, 3.5-fold; and ACACB, 3.2-fold), and fatty acid synthase (FASN, 8.5-fold increase) (Figs 1D, 1F, Table S1). This is consistent with previous results indicating that *de novo* lipogenesis is a central feature of thermogenesis in BAT^15,16^.UCP1 increases 3.7-fold, and the mitochondrial pyruvate carrier 2 (MPC2) increases 3.1-fold after mild cold acclimation (Fig S1A, Table S1). All of the proteins mentioned above also significantly increase after severe cold acclimation. In addition, proteins related to the electron transport chain (ETC) complex I are among the most upregulated after severe cold acclimation (NDUFA12, increasing 3.7-fold; NDUFA10, 2.4-fold; NDUFB6, 3.7-fold; and MTND3, 4.4-fold) (Figs 1F, S1A, Table S1). Interestingly, the calcium load-activated channel TMCO1 increases 4.6-fold after severe cold acclimation, but is not reliably measured in the mild cold condition, possibly because it is below the limit of detection (Figs 1E, S1A, Table S1). Notable decreasing proteins include the selenium binding protein 1 (SELENBP1, 2.9-fold decrease after severe cold acclimation), Mimecan (OGN, 4.7-fold), and the centromere protein V (CENPV, 3.5-fold), all of which also significantly decreased, albeit with lower fold-changes, after mild cold acclimation (Fig 1D, 1E, S1B, Table S1). Interestingly, some proteins decrease after mild cold acclimation, and increase to near thermoneutral levels in the severe cold. These include the long-chain fatty acid CoA ligase (ACSL5, 4.8-fold decrease after mild cold acclimation), and the ubiquitin-like modifier SUMO2 (6.3-fold decrease) (Figs 1D, S1B, Table S1). Taken together, analysis of the most significant cold-induced protein changes confirms that lipid synthesis and electron transport are simultaneously upregulated during BAT thermogenesis, while revealing other novel, potential protein markers of BAT activity.

Despite not being among those most significantly regulated, many other notable proteins show interesting responses to cold acclimation. For example, the insulin-regulated glucose transporter, SLC2A4 (a.k.a. GLUT4), is upregulated 2.7-fold in mild cold consistent with insulin and beta-adrenergic signaling cooperating during CIT (Table S1). Early glycolytic enzymes are also significantly cold-induced, including hexokinase 2 (HK2, 2.6-fold increase in severe cold), glucose-6 phosphate isomerase (GPI, 2.9-fold increase), and the phosphofructokinase liver isoform (PFKL, 2.7-fold increase) (Table S1). This is consistent with increased glucose flux into active BAT and is the basis for detecting BAT by FDG-PET-CT imaging^2^. Enzymes in the pentose phosphate pathway are also significantly induced, including glucose-6-phosphate 1-dehydrogenase X (G6PDX, 2.7-fold increase in severe cold), ribose-5-phosphate isomerase (RPIA, 2.6-fold increase, q = 0.14), and transaldolase (TALDO1, 1.8-fold increase), suggesting important destinations for glucose besides pyruvate synthesis (Fig S1A, Table S1). In addition, the carnitine palmitoyltransferase enzymes CPT1B and CPT2 increase after severe cold acclimation, 2-fold and 1.6-fold respectively (Table S1), suggesting an increase in fatty acid import into mitochondria, which is thought to be rate-limiting for fatty acid oxidation. Thus, manual inspection of proteins related to metabolism provides more detailed insight about glucose and fatty acid metabolism during BAT thermogenesis.

To further characterize the thermogenic proteome, we performed a gene ontology (GO) enrichment analysis for proteins that increase or decrease in *either* mild *or* severe cold. GO Process analysis identifies “mitochondrial ATP synthesis-coupled electron transport”, “glucose 6-phosphate metabolic process”, “acyl-CoA metabolic process”, and “fatty acid biosynthetic process” as being enriched among proteins that increase in mild or severe cold relative to thermoneutrality (Fig 1G). Analysis of GO Components for these upregulated proteins shows enrichment of the terms “mitochondrial respiratory chain complex I” and “NADH dehydrogenase complex”, suggesting an overall increase in complex-I proteins (Fig 1G). While there are several significantly decreasing proteins, named above, we found that no corresponding GO pathways are enriched. Together, our global proteomics analysis of BAT is consistent with cold acclimation stimulating glucose uptake, and flux towards glycolysis, the pentose phosphate pathway, *de novo* lipogenesis, capacity for fatty acid oxidation, and electron transport.

### Mitochondrial proteome remodeling during thermogenesis

Because BAT thermogenesis increases mitochondrial metabolic activity, we used our proteomics dataset to characterize the mitochondrial proteome in response to the different cold acclimation conditions. First, we annotated all measured proteins as either “mitochondrial” or “non-mitochondrial” based on GO Component annotations, and compared the abundance of the median mitochondrial protein to the median non-mitochondrial protein across all replicates and conditions. This analysis revealed that mitochondrial proteins significantly increase relative to non-mitochondrial proteins after both mild and severe cold acclimation (Fig 2A). We also examined the relative changes to proteins annotated to different mitochondrial subcompartments. This approach indicates a greater increase in inner mitochondrial membrane proteins, relative to mitochondrial matrix proteins, in cold conditions relative to thermoneutrality (Fig 2B). This suggests that mitochondria in BAT contain more dense cristae after cold acclimation. We also observe a similar, but milder, increase in outer mitochondrial membrane proteins relative to mitochondrial matrix proteins (Fig 2B), suggesting increased mitochondrial number along with decreased size of each mitochondrion.

**Figure 2:**
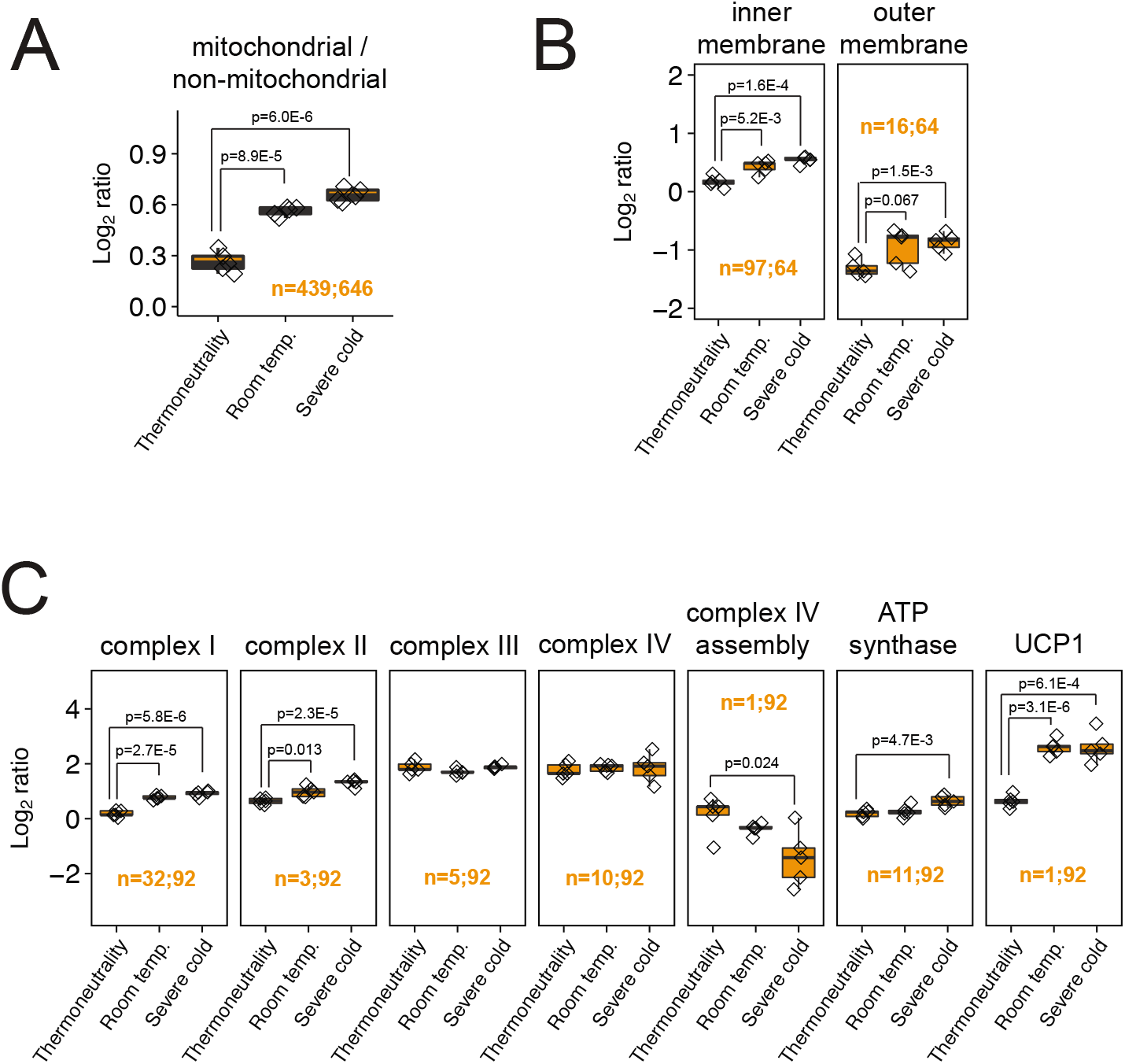
Mitochondrial proteins as a whole, and ETC complex-I proteins in particular, increase their relative abundance during thermogenesis. A. Plot showing changes to the Log_2_ ratio of the mean mitochondrial to the mean non-mitochondrial protein. The number of proteins used to calculate medians for mitochondrial and non-mitochondrial fractions are indicated for each experiment before and after the semicolon, respectively. Data points represent ratios calculated from individual animals. B. Plot showing the Log2 ratio of the mean inner membrane or outer membrane protein to the mean mitochondrial matrix protein. The number of inner/outer membrane proteins and the number of matrix proteins are indicated for each experiment before and after the semicolon, respectively. Data points represent ratios calculated from individual animals. C. Plot showing the Log_2_ ratio of the mean ETC complex protein to the mean mitochondrial matrix protein. The number of ETC complex proteins and the number of matrix proteins are indicated for each experiment before and after the semicolon, respectively. Data points represent ratios calculated from individual animals.

To further investigate changes to inner mitochondrial membrane proteins, we calculated the ratio of each complex in the ETC to the mitochondrial matrix, using annotation from WikiPathways to define proteins that represent complexes I-IV and the ATP synthase complex. Strikingly, complex I and complex II proteins, but not those from complexes III and IV, increase in response to all cold acclimation conditions (Fig 2C), and show a nearly two-fold enrichment in mitochondria between thermoneutrality and severe cold adaptation. The ATP synthase complex also has a mild cold-dependent increase, which could reflect a compensatory mechanism to generate sufficient ATP in the face of uncoupled respiration (Fig 2C). These results suggest that the stoichiometry of ETC complexes, specifically complexes I and II, is regulated during BAT thermogenesis.

### Overlap between the cold-induced transcriptome and proteome

Compared to the thermogenic proteome, the RNA-seq transcriptome analyses of these BAT samples, reported in^15^, contains a similar distribution of fold-changes between conditions (Fig 3A), but a greater percentage of significant changes at a 5% FDR threshold (Fig 3B). 88% of proteins that significantly increase (q ≤ 0.05) in room temperature acclimation, and 68% of those that significantly increase in severe cold acclimation, also increase at the mRNA level (Fig 3C). Among the overlapping mRNA-protein pairs that most significantly increase in cold in both the proteomic and transcriptomic analyses are fatty acid synthesis enzymes (including ACLY, ACSS2, ACACA, ACACB, FASN, SCD1); AGPAT2, which functions in triacylglyceride synthesis; and UCP1 (Fig 3D). Correlation between mRNA-protein pairs is less consistent in transcripts or proteins that decrease after mild or severe cold acclimation (Fig 3C), which may suggest regulation via protein translation or degradation. Indeed, a recent study found that proteasomal activity increases in cold-adapted BAT, and is important for effective thermogenesis^17^. Unlike at the protein level, there was no increase in transcripts encoding for inner or outer mitochondrial membrane, or ETC proteins, relative to mitochondrial matrix proteins after cold acclimation (Fig 3E). This suggests that regulation of the mitochondrial proteome with BAT activation is post-translational.

**Figure 3:**
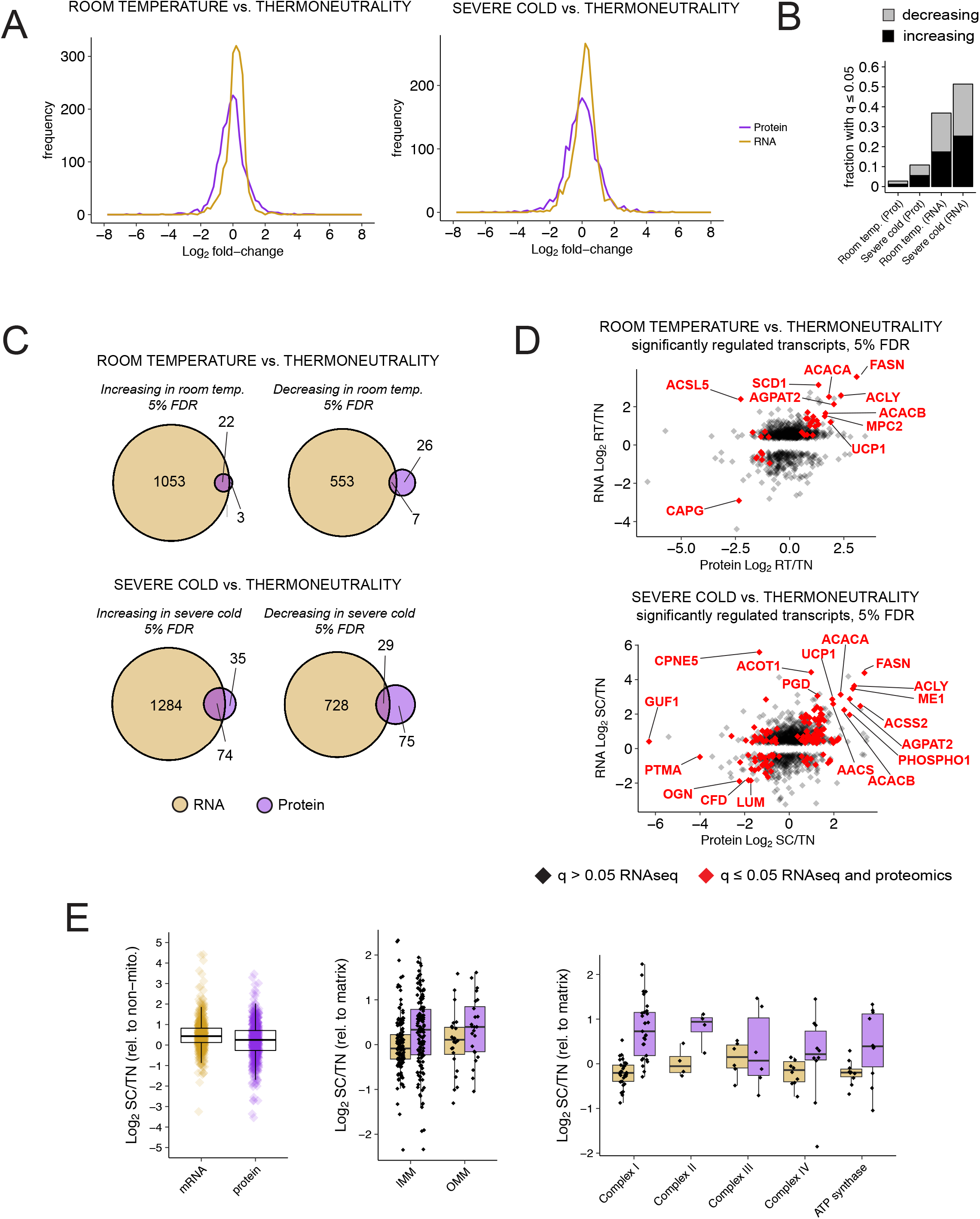
Comparison of proteome and transcriptome profiling of 3-temperature cold acclimated mouse BAT. Transcriptome profiling from Sanchez-Gurmaches et al.^15^ was compared to the 3-temp proteomics experiment presented in this study. A. Distribution of Log_2_ fold-changes, both room temperature (top) and severe cold (bottom) vs. thermoneutrality, for the proteome (purple) and transcriptome (gold) profiling datasets. B. FDR q-values were compared between each condition and the thermoneutrality control. The fraction of proteins or transcripts with q ≤ 0.05 is plotted for the indicated experimental conditions. C. Euler plots comparing significantly (q ≤ 0.05) upregulated or downregulated proteins and their respective transcripts. Only transcripts whose proteins were detected in the proteomics experiment are shown. D. Transcripts that increase after room temperature (top) or severe cold (bottom) acclimation are plotted to compare fold-changes between the transcriptome and proteome datasets. Sites that are significantly (q ≤ 0.05) regulated in both experiments are shown in red, even if their direction of change differs. E. Analysis of mRNA and protein levels in mitochondrial sub-groups. Left: the ratio of mitochondrial to non-mitochondrial mRNA/protein). Middle: the ratio of inner mitochondrial membrane (IMM) or outer mitochondrial membrane (OMM) to mitochondrial matrix mRNA/protein. Right: the ratio of ETC complexes to mitochondrial matrix mRNA/protein.

### Mitochondrial protein acetylation increases during cold acclimation

Because multiple routes to acetyl-CoA generation are increased in active BAT, we hypothesized that cold adaptation is also associated with significant changes in protein acetylation. To test how BAT activation affects protein acetylation patterns, we digested total BAT protein with trypsin, enriched for acetylated peptides using anti-acetyl-lysine antibodies, then quantified their amount using label-free mass spectrometry. Across all conditions, the number of acetylation sites, or acetylsites, identified per run ranged from 794 to 1904, with higher number of identified acetylation sites in the severe cold condition (Fig 4A). To account for the fact that acetylated proteins might themselves change in abundance between conditions, acetylsite quantifications were normalized to the level of the respective protein for that replicate. After this normalization, we calculated q-values for room temperature and severe cold conditions relative to thermoneutrality (Table S2).

**Figure 4:**
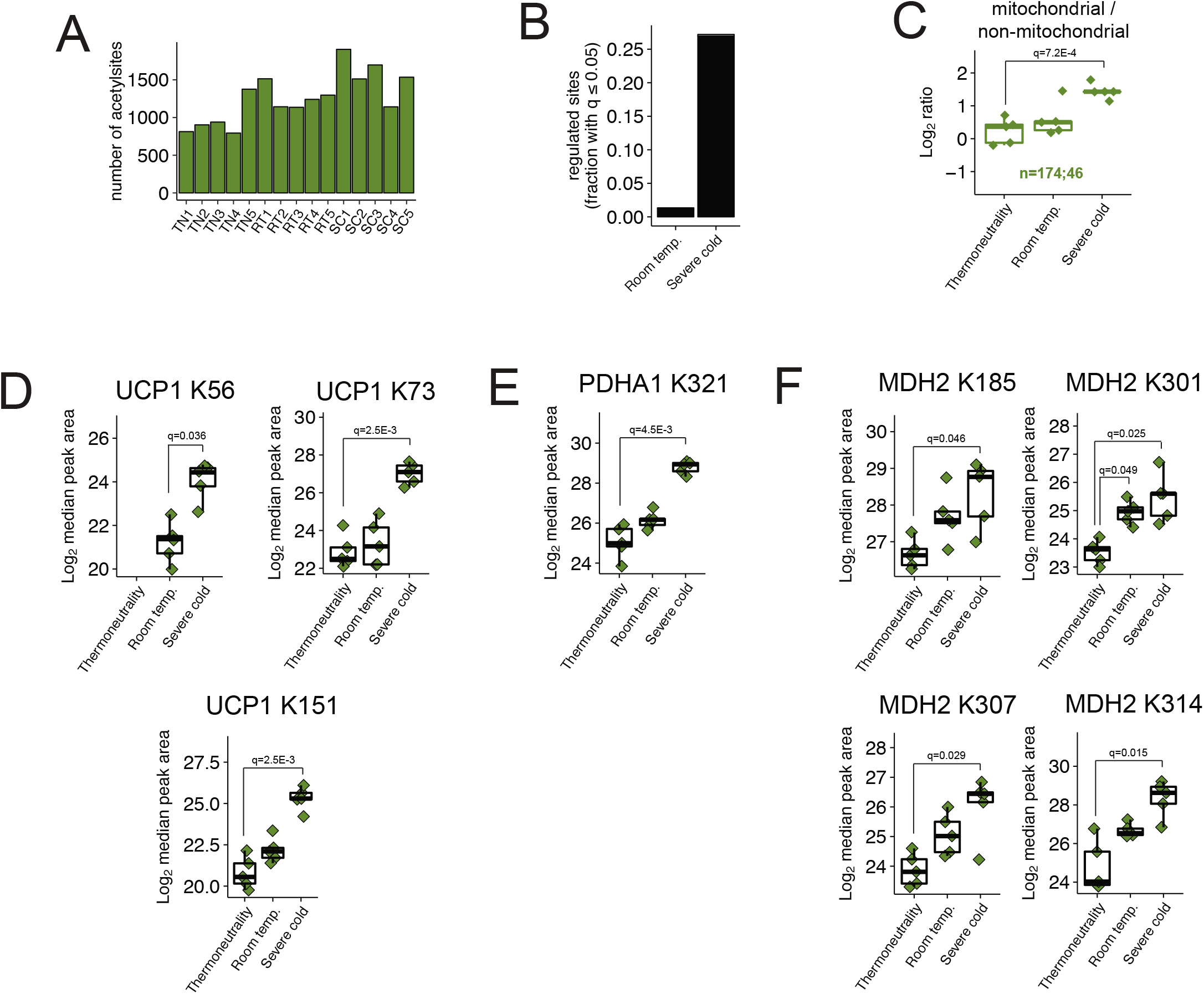
Mitochondrial protein acetylation increases during thermogenesis. A. Number of identified acetylsites in each biological condition. Error bars represent standard deviation. B. The fraction of acetylsites with q ≤ 0.05, either increasing (black) or decreasing (grey), is plotted for the indicated experimental conditions, relative to the thermoneutrality control. C. Log_2_ ratios of the mean mitochondrial to mean non-mitochondrial protein-normalized acetylated peptide forms, across all conditions and experiments. The number of mitochondrial and non-mitochondrial proteins is indicated for each experiment before and after the semicolon, respectively. Data points represent ratios calculated from individual animals. D. Acetylation on UCP1 K56, K73, and K151 and is shown for all three temperature conditions. Data points represent measurements from individual animals. E. Acetylation on PDHA1 K321 is shown for all three temperature conditions. Data points represent measurements from individual animals. F. Acetylation on four MDH2 sites is shown for all three temperature conditions. Data points represent measurements from individual animals.

Cold-induced thermogenesis caused a widespread increase in protein acetylation in BAT. At 5% FDR, over 25% of protein-normalized acetylation sites increase in severe cold acclimation, while only 1% increase in room temperature acclimation (Fig 4B). The vast majority of acetylation changes occur on mitochondrial proteins. GO enrichment analysis on acetylsites that increase after severe cold acclimation reveals terms related to the mitochondria and fatty acid catabolism (Figs S2A, S2B). Acetylation on mitochondrial proteins, relative to non-mitochondrial proteins, increases over 2-fold in response to severe cold acclimation (Fig 4C).

Interestingly, UCP1 is hyperacetylated after cold acclimation on K151 (22.6-fold increase after severe cold), K73 (17.7-fold increase), and K56 (7.2-fold increase from mild to severe cold) (Fig 4D, Table S2). UCP1 localizes to the inner mitochondrial membrane, and all three of these sites are on the mitochondrial matrix-facing side. Several other hyperacetylated mitochondrial proteins are particularly noteworthy based on the number of acetylsites that significantly increase (Table S2). In the TCA cycle, two isocitrate dehydrogenase isoforms are highly hyperacetylated in cold acclimation; IDH2, with 12 acetylsites, and IDH3 alpha subunit, with 8 acetylsites; the alpha subunit of the fatty acid oxidation trifunctional enzyme, HADHA, is hyperacetylated at 12 sites; and dihydrolipoyl dehydrogenase (DLD), which functions in branch chain amino acid catabolism, is hyperacetylated at 8 sites.

Relative to mitochondrial protein acetylation, we detected few changes in cytoplasmic protein acetylation (Figs S2C, S2D). Histone H3 acetylation regulates gene expression; however, we did not detect significant changes to histone H3 acetylation relative to its total protein levels (Figs S2E, S2F). Because of low nuclear histone abundance, we speculate that a higher resolution analysis of histone marks at specific promoters via techniques such as ChIP-seq is required to fully assess acetylation-mediated epigenetic changes that might be associated with thermogenesis. These findings suggest that the nuclear-cytoplasmic acetyl-CoA generated in active BAT may predominantly be used for *de novo* lipid synthesis rather than protein lysine acetylation, and that the increase in mitochondrial protein acetylation relative to other compartments may be driven by the robust increase in mitochondrial nutrient (e.g. fatty acid, pyruvate) oxidation also associated with thermogenesis.

We also identified cold-induced acetylation sites on mitochondrial proteins with previously characterized metabolic functions. For example, acetylation of the pyruvate dehydrogenase E1 alpha subunit (PDHA1) on K321 has been shown to be inhibitory, and implicated in the Warburg effect in cancer cells^18^. PDHA1 K321 acetylation increases in severe-cold-acclimated BAT (Fig 4E, Table S2). Inhibiting PDHA1 in BAT could thus temper the flux from glycolysis to acetyl-CoA synthesis, and favor other uses of glycolytic intermediates. Another example of hyperacetylation with previously characterized function is on the the mitochondrial enzyme malate dehydrogenase (MDH2). MDH2 functions in the TCA cycle and as part of the malate-aspartate shuttle that transports NADH reducing equivalents between the mitochondria and cytosol. MDH2 has four acetylation sites that, when simultaneously mutated to arginine, decrease its activity^19^. We find that all four of these acetylsites on MDH2 significantly increasing in severe-cold (Fig 4F, Table S2). The aforementioned cold-induced acetylation sites on PDHA1 and MDH2 could be be important for regulating glucose and NADH metabolism, respectively, in the context of BAT thermogenesis.

### Analysis of independently temperature-acclimated mice confirms key proteomics and acetylproteomics findings

To solidify our findings, we repeated our BAT proteome and acetylproteome analysis on an independent mouse cohort. This second experiment was largely similar to the previous one with regards to animal age, diet, and housing conditions (see Methods), but here we increased the number of replicates (7 mice for room temperature, 6 mice for thermoneutrality) and focused on the two temperatures that are relevant to humans^20,21^: thermoneutrality and room temperature.

We first compared the thermogenic proteome datasets and find they are of similar scope and quality as they contain a similar number of identified proteins (Fig S3A), and have a similar degree of quantitative reproducibility indicated by the distribution of coefficients of variation (Fig S3B). The percentage of significantly regulated proteins is greater in the second experiment (Fig S3C); and the magnitude and directionality of proteins that are significantly altered in at least one experiment are in good agreement (Fig S3E). Thirteen proteins significantly increase in both experiments, including *de novo* lipogenesis enzymes ACACA, ACACB, and FASN, and the early glycolytic enzymes HK2 and GPI (Tables S1, S3). Relevant proteins that significantly increased in one dataset (e.g. ACLY, ACSS2, CPT1B and CPT2), also show an increase in the other dataset (Tables S1, S3). We found 2 proteins significantly decreasing in both datasets: the extracellular matrix protein FBN2 and the cytosolic selenium-binding protein SELENBP1 (Fig S3D, Tables S1, S3). The cold-stimulated increase in mitochondrial proteins relative to non-mitochondrial proteins is also conserved between both data sets (Fig S3G), as is an increase in the amount of outer mitochondrial membrane proteins (Fig S3H) and complex I proteins (Fig S3I) relative to mitochondrial matrix proteins.

Mitochondrial protein hyperacetylation occurs during mild cold acclimation in the second experiment, to an even greater extent than in the first experiment. Importantly, the summary statistics for acetylproteome measurements are comparable between the two datasets, both in terms of the number of identified sites (Fig S4A) and the quantitative reproducibility (Fig S4B). While the number of significantly regulated sites is greater in the second experiment (Fig S4C), due in part to to increased replicates, the global increase in mitochondrial protein acetylation over cytoplasmic/nuclear protein acetylation is consistent (Fig S4D). Specific acetylation sites of interest, discussed above–PDHA1 K321; MDH2 K185, K301, K307 and K314–all significantly increase in room temperature relative to thermoneutrality acclimation in the second experiment (Fig S4E-G, Table S4). Expression of UCP1 between chronic thermoneutral and room temperature housing showed no differences in this experiment (Fig S5A, Table S3), yet the significant increase in acetylation on UCP1 K151 was conserved (Fig S5B, Table S4). The second experiment also revealed a 13.6-fold increase in acetylation on UCP1 K67 after mild cold acclimation (Fig S5B, Table S4) that did not pass our significance threshold in the first experiment. Taken together, these data add confidence to our proteomics and acetylproteomic findings and solidify many of our hypothesis on how thermogenesis regulates BAT metabolism.

### Cold acclimation alters glycolytic intermediates, nucleotide triphosphates, and electron transporters in BAT

Next we explored the BAT metabolic landscape by measuring 124 polar metabolites extracted from BAT by LC-MS/MS. Measured metabolites include intermediates to glycolysis, the TCA cycle, and amino acid and nucleotide metabolism. 23 metabolites increase in abundance in severe-cold-adapted BAT relative to thermoneutrality at 5% FDR while 13 metabolites decrease. Cyclic AMP is among those increasing with cold acclimation consistent with its role in transducing the beta-adrenergic stimulus to protein kinase A (PKA) during thermogenesis (Figs 5A, S6A, Table S5). The nucleotide triphosphates ATP, GTP, and UTP all decrease significantly at least 6-fold in cold-adapted BAT (Figs 5A, S6B, Table S5), which is consistent with the decreased oxidative phosphorylation that accompanies uncoupled respiration^22^. The corresponding loss of nucleotide triphosphates other than ATP would be an expected equilibrium state due to the action of nucleotide-diphosphate kinase^23^.

**Figure 5:**
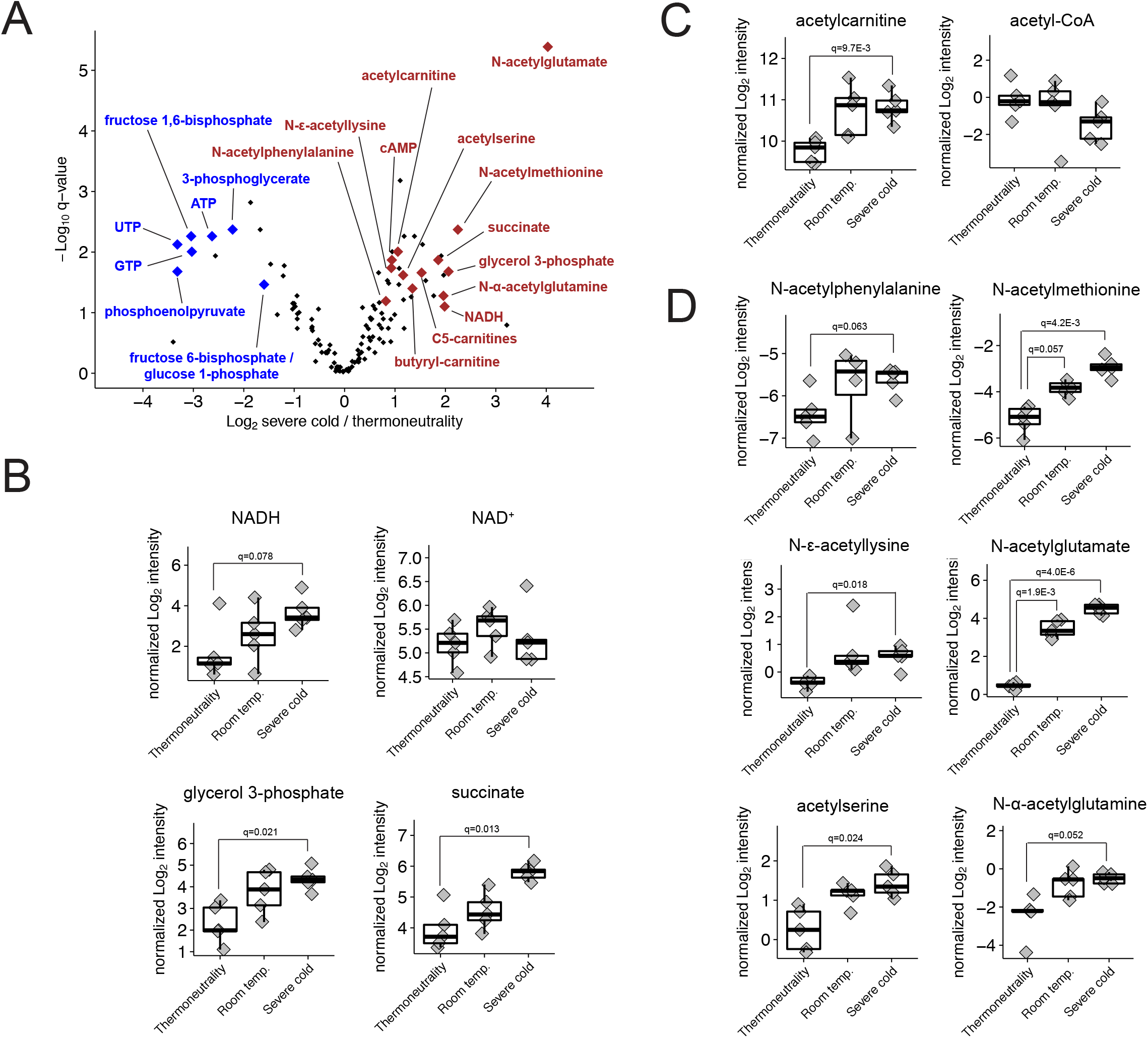
Polar metabolite measurements reveal many changes to acetylated metabolites and reducing equivalents. A. Volcano plot showing the Log_2_ fold-change versus the -Log_10_ q-value for all metabolites, between the severe cold and thermoneutrality conditions. Selected increasing (brown) and decreasing (blue) metabolites are indicated. B. Barplots showing quantifications for metabolites involved with cellular redox reactions. Data points represent measurements from individual animals. C. Barplots showing quantifications for acetylcarnitine and acetyl-CoA. Data points represent measurements from individual animals. D. Barplots showing quantifications for acetylated amino acids. Data points represent measurements from individual animals.

Several glycolytic intermediates decrease with cold-acclimation including fructose 1,6-bisphosphate, 3-phosphoglycerate, phosphoenolpyruvate, and a saccharide that could be either fructose-6-phosphate (glycolysis) or glucose-1-phosphate (glycogen metabolism) (Figs 5A, S6C, Table S5). Decreased steady-state levels of glycolytic metabolites is consistent with increased glycolytic flux. In contrast, glycerol 3-phosphate increases 4.2-fold with severe cold acclimation (Fig 5B, Table S5). Glycerol 3-phosphate synthesis occurs by several mechanisms, including via glycolysis, and is a precursor to triglyceride synthesis. In rats, glucose-derived glyceride levels in BAT increased after the tissue was activated by a high-fat cafeteria diet^24^. Thus, it is possible that synthesis of glycerol 3-phosphate and its subsequent incorporation into triglycerides is also an important role for glucose in cold-activated BAT. Another possibility is that glycerol 3-phosphate increases by hydrolysis of triglycerides due to the increased lipolysis rate. In addition to being a precursor to triglycerides, glycerol 3-phosphate is also an electron carrier that can transfer reducing equivalents between the mitochondria and cytosol, where it can be consumed in a reaction, along with NAD^+^, to generate NADH and dihydroxyacetone phosphate. Interestingly, two other reduced electron carriers increase after cold acclimation: succinate and NADH. Succinate increases 3.6-fold in the severe cold (Figs 5A, 5F, Table S5), while NADH increases 3.9-fold with q = 0.078 (Fig 5B, Table S5). NAD^+^ does not change in response to cold acclimation (Fig 5B, Table S5). This is interesting, given the importance of reducing equivalents in driving the ETC, as well as their potential for participating in product inhibition of the enzymes that generate them. In sum, these results (summarized in Fig 6) indicate significant changes to glycolysis, nucleotide triphosphates, and redox biochemistry in BAT following cold acclimation.

**Figure 6:**
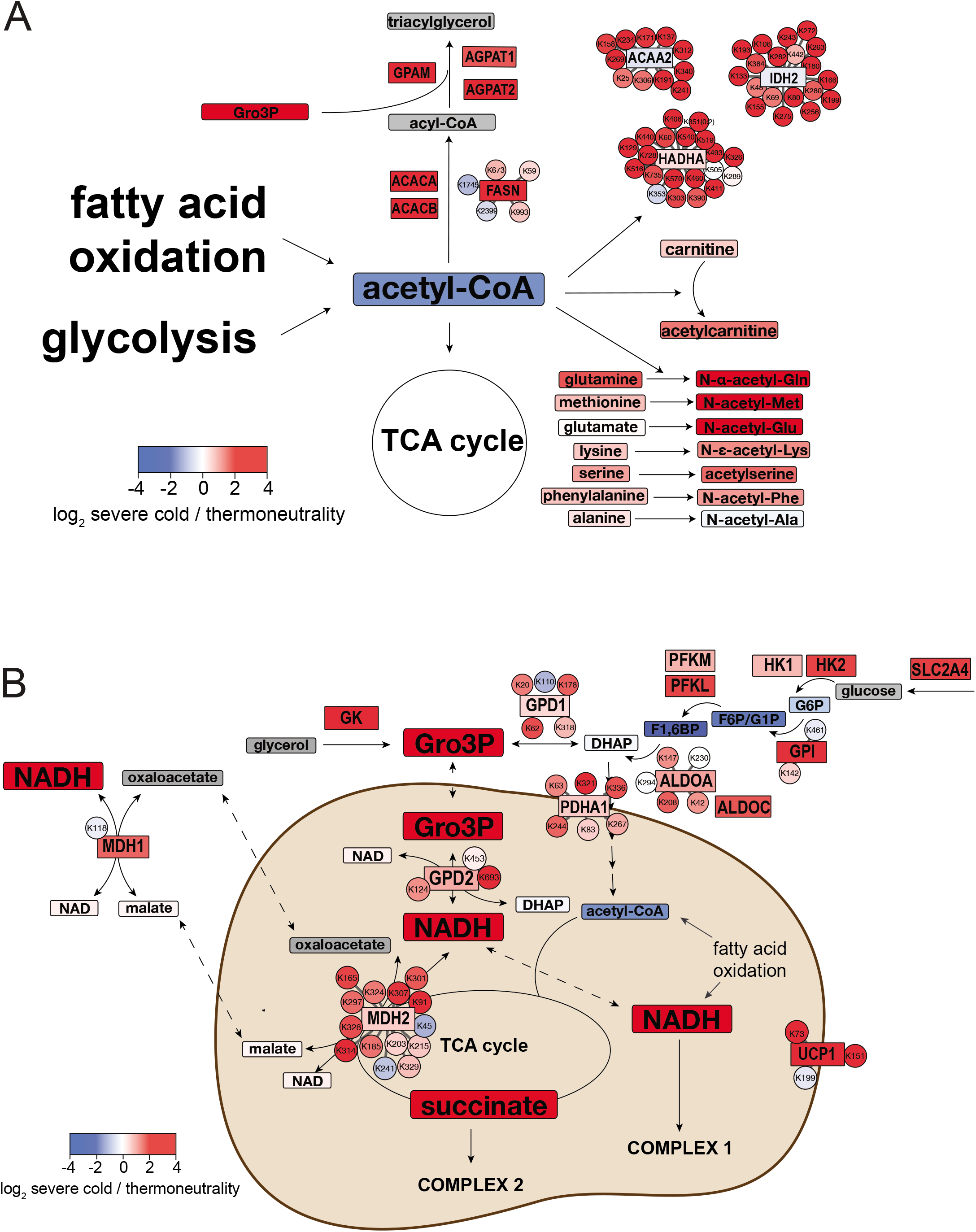
Chronic cold acclimation alters acetyl-CoA usage and redox metabolism in BAT. A. Schematic showing quantitative data for select metabolites, proteins, and acetylation sites for selected pathways that consume acetyl-CoA. B. Schematic showing quantitative data for select metabolites, proteins, and acetylation sites that are connected to cellular redox.

### Increases to acetylated metabolites during thermogenesis suggests altered acetyl-CoA flux

Acetyl-CoA levels do not change between the thermoneutral and room temperature conditions and mildly decrease with severe cold (Fig 5C, 6A, Table S5). Thus, while the rate of acetyl-CoA synthesis would be expected to rise with cold due to the increase in fatty acid oxidation and *de novo* lipogenesis that is characteristic of active BAT, the constant or decreasing acetyl-CoA abundance likely reflects increased acetyl-CoA flux towards protein acetylation and perhaps other metabolites. Acetyl-CoA is a highly reactive metabolic intermediate due to the high energy thioester bond linking its acetyl group to the CoA moiety, and there are several cellular reactions that can consume it (Fig 6A); thus its availability must be under tight control. Indeed, we find increased acetylation of acetylcarnitine and acetylated amino acids in thermogenic BAT. Acetylcarnitine is generated in the mitochondria from acetyl-CoA and carnitine by the enzyme carnitine acetyltransferase, and exported from the mitochondria to buffer against high mitochondrial acetyl-CoA levels^25^. Acetylcarnitine is upregulated 2.1-fold in severe-cold-acclimated BAT, suggesting mitochondrial acetyl-CoA synthesis may exceed its utilization by the TCA cycle in cold-stimulated BAT (Fig 5C, Table S5). Interestingly, we also find that four acetylated amino acids also significantly increase in abundance with cold: N-acetylglutamate (16-fold increase), N-acetylmethionine (4.8-fold), acetylserine (2.2-fold), and N-ε-acetyllysine (1.9-fold) (Figs 5D, 6A, Table S5). Two more acetylated amino acids (acetylphenylalanine and N-α-acetylglutamine) increase by greater than 2-fold (Fig 5D, Table S5).

BAT thermogenesis may lead to accumulation of some metabolites upstream of acetyl-CoA as well, as the branched chain amino acids valine, leucine, and isoleucine, which are catabolized to acetyl-CoA, all increase with cold (Table S5). In addition, amino acids could be generated by protein degradation, which has been shown to be increased in active BAT^17^. While we do not measure subcellular fractions of these metabolites, correlation with acetylcarnitine and mitochondrial protein acetylation levels suggest a role for acetylated amino acids in buffering mitochondrial acetyl-CoA.

### Functional analysis of UCP1 acetylation sites

Finally, given the BAT specificity and function of UCP1 in thermogenesis, and the cold-induced regulation observed for its four acetylation sites detected, we wondered if these sites might be functionally relevant. When normalized to protein levels, acetylation on K56, K67, K73, and K151 increase in response to either mild or severe cold, with q ≤ 0.05 in either of the two proteomics experiments (Figs 4D, S5B). To test potential functional relevance of UCP1 acetylation on these sites, we mutated K56, K67, K73 and K151 to either arginine (UCP1-4KR), which blocks acetylation, or glutamine (UCP1-4KQ), which mimics acetylation. We also generated UCP1-3KR and UCP1-3KQ by only mutating K56, K67 and K73 (Fig 7A). We first expressed recombinant UCP1 mutant constructs and a wild-type UCP1 (UCP1-WT) control in 293T cells. Western blot analysis indicated that the 3KR and 4KR mutants express at similar levels to the UCP1-WT control. However, both UCP1-3KQ, and more dramatically UCP1-4KQ, express at significantly lower levels (Fig 7B). This is not due to differences in mRNA expression because UCP1-4KQ and -3KQ show higher transcription than control (Fig 7C). To test whether acetylation alters UCP1 stability, we treated the 293T cells expressing UCP1-WT, -4KR and -4KQ with cycloheximide (CHX) to inhibit *de novo* protein synthesis and estimate UCP1 protein halflives. We observe that the acetyl-mimetic mutant (UCP1-4KQ) has a shorter half-life relative to UCP1-WT, while UCP1-4KR has a longer half life (Fig 7D). While the exact mechanism of how acetylation may influence UCP1 stability remains to be determined, and limitations of overexpressing wild-type and mutant UCP1 *in vitro* are duly noted, our results show preliminary evidence that cold-induced acetylation of UCP1 could have functional relevance.

**Figure 7:**
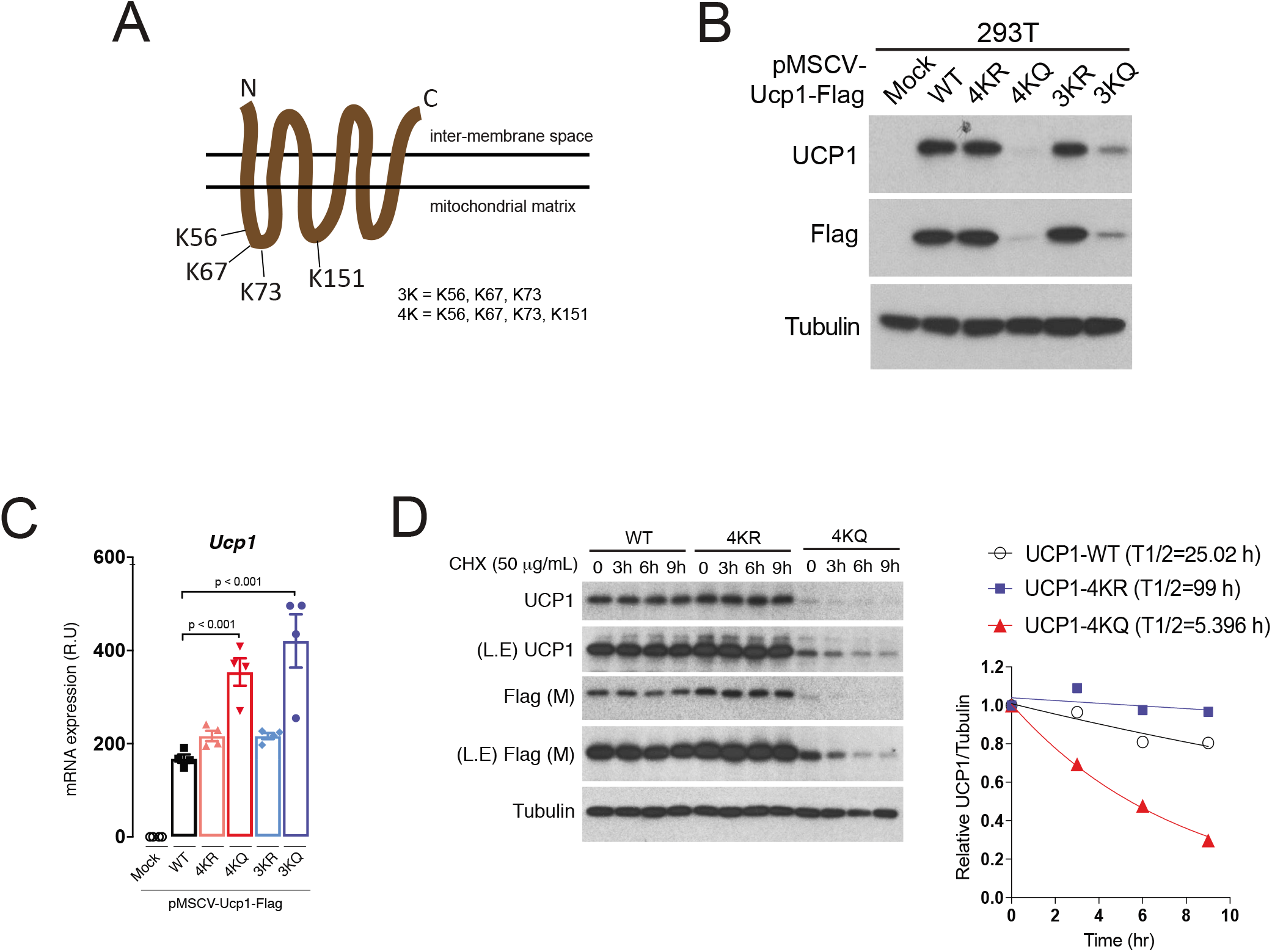
Four cold-dependently hyperacetylated lysine residues on UCP1 are important for stability. A. Outline of UCP1 acetylation mutant constructs design. Note that all four acetylation sites are predicted to be mitochondrial matrix-facing area. B. Immunoblots of UCP1 wild-type or mutant constructs expressed in 293T cells. C. mRNA expression levels, measured by qPCR, of UCP1 wild type or mutant constructs expressed in 293T cells. Data points are biological replicates, and error bars represent standard error of the mean. D. Immunoblot (left) and quantification (right) showing protein expression of UCP1 wild-type or mutant constructs in a time series after cycloheximide (CHX) treatment. Half-lives of UCP1 proteins are calculated in the graph. In all immunoblot analysis, expression of alpha-tubulin was used as a loading control.

## DISCUSSION

Brown adipose tissue is of great interest as a potential therapeutic target for type-2 diabetes and other metabolic diseases associated with obesity, yet the molecular underpinnings of BAT metabolic activity have not been fully elucidated. Here, we sought a better understanding of the BAT molecular landscape by comparing proteome, acetylproteome, and metabolome data between inactive and different degrees of cold-activated BAT. We show that cold-induced thermogenesis has significant and reproducible effects on all three “omics” measurements taken in this study. Among many findings, our integrative analysis suggests a broad reprogramming of acetyl-CoA metabolism during thermogenesis. This conclusion is supported by (1) the upregulation of enzymes related to *de novo* lipogenesis, which consumes cytosolic acetyl-CoA, (2) increases to carnitine palmitoyltransferases CPT1B and CPT2, suggesting increased capacity for fatty acid oxidation, which generates mitochondrial acetyl-CoA, (3) the upregulation of mitochondrial protein acetylation, and (4) increases in acetylated metabolites, including acetylcarnitine as well as several acetylated amino acids.

Many mitochondrial proteins were hyperacetylated in BAT after cold acclimation, including a few sites on metabolic enzymes that have previously-characterized functions. We find that PDHA1 K321 becomes hyperacetylated after cold-acclimation relative to PDHA1 protein levels (Fig S4E). PDHA1 acetylation at this site has been shown to promote a transformed phenotype in cancer cell lines, likely via the Warburg effect^26^. A Warburg-like effect could exist in cold-activated BAT, whereby glycolytic intermediates could be directed to other key metabolites. Another example of hyperacetylation at previously-characterized sites is on MDH2. We report that several lysine residues on the mitochondrial malate dehydrogenase, MDH2, are hyperacetylated in cold-adapted BAT (Fig 4F). Some of these sites have been shown to increase the activity of the MDH2 reverse reaction, which consumes NADH and generates malate, which is shuttled out of the mitochondria^19^. This could be a mechanism by which NADH product inhibition is overcome in active BAT (Fig 6B). Finally, we found cold-dependent hyperacetylation on four sites on UCP1, and through mutagenesis experiments and cell culture expression, we demonstrate increased degradation of UCP1 with acetylation-mimicking mutations, suggesting that acetylation at these residues may influence the stability of UCP1 and be functionally relevant.

BAT thermogenesis is centered around the ETC. Our metabolomics data revealed cold-dependent increases to three ETC substrates: NADH, succinate, and glycerol-3 phosphate (Fig 6B). This could reflect metabolic reprogramming geared toward generating substrates for electron transport. In addition, succinate was recently shown to be sequestered from circulation in BAT, where it subsequently potentiates thermogenesis via reactive oxygen species (ROS) generation by complex II^27^. Alongside increases in substrates for electron transport, we observe changes to ETC complex stoichiometry that could also affect the generation of ROS. ETC complexes have been shown to assemble in cells as supercomplexes^28^. One study in neurons and astrocytes showed that adjusting expression of a complex-I protein affects the presence of complex-I in a free state versus in supercomplexes, which in turn affects the production of ROS^29^. ROS production increases in BAT during cold acclimation, and has been shown to be necessary for complete induction of thermogenesis^30^, but our knowledge of the mechanisms of ROS production is incomplete. We observe increases to complexes I-II of the ETC, and not other complexes, which could be a way to adjust ETC stoichiometry and consequent production of ROS during thermogenesis.

Turning to the observed acetyl-CoA reprogramming in thermogenesis, another mechanism by which cells regulate mitochondrial acetyl-CoA levels is through the action of carnitine acetyl-transferase (CRAT). One recent study showed that deletion of CRAT in skeletal muscle leads to an increase in acetyl-CoA and a concomitant increase in mitochondrial protein acetylation in response to high-fat feeding^31^. In our study, acetylcarnitine levels increase as a result of mild and severe cold adaptation suggesting a key role for CRAT in buffering mitochondrial acetyl-CoA pools during BAT thermogenesis. Like acetylcarnitine, acetylated amino acids increase in BAT after cold acclimation. Nobody has reported amino acid acetylation as a means to dispense of excess acetyl-CoA, but it is a plausible mechanism that could work in parallel with acetylcarnitine synthesis. While it is also possible that the increase in alpha-amino acetylated amino acids is the result of increased cleavage of N-terminally acetylated protein residues, we found no evidence of increased N-terminal acetylation in the proteomics data (Fig S7). Thus, we postulate that conversion to citrate and subsequent cytosolic *de novo* lipogenesis, as well as the acetylation of amino acids, carnitine, and lysine residues on mitochondrial proteins, function to consume mitochondrial acetyl-CoA generated from glucose and fatty acids during BAT thermogenesis (Fig 6A). This would allow oxidative metabolism to continue with minimized product inhibition by acetyl-CoA. Interestingly, the most upregulated metabolite in our dataset is N-acetylglutamate, which stimulates urea cycle activity in the liver by allosterically activating the enzyme carbamoyl phosphate synthetase^32^. Given that proteasomal activity is known to increase in active BAT^17^, increased urea cycle activity may be necessary to dispose of nitrogen by products from amino acid catabolism. In the future, it will be interesting to measure acetyl-CoA in distinct subcellular fractions (e.g. mitochondrial, nuclear, and cytosolic), as well as flux of reactions generating and consuming acetyl-CoA in order to characterize the connection between acetyl-CoA and acetylated amino acids. In addtion, the dramatic increase of N-acetylglutamate suggests that simultaneous, multi-omics analysis of multiple tissues could help to elucidate novel inter-organ crosstalk mechanisms during thermogenesis.

In conclusion, we demonstrate that cold-induced thermogenesis profoundly alters the BAT proteome by increasing the levels and/or the acetylation of specific proteins. We also describe many changes in key metabolites in response to cold-induced thermogenesis that provide important clues to how BAT fuels its metabolism. This work emphasizes the metabolic flexibility of BAT and has important implications for development of therapeutic strategies targeting BAT metabolism to fight metabolic diseases.

## ACKNOWLEDGMENTS

We would like to thank all members of the Villen lab for critical discussions and feedback on the project. This work was supported by NIH grants R01 CA196986 (to D.A.G.); R35 GM119536, R01 AG056359, and R01 NS098329 (to J.V.); R37 DK030898 and R01 DK103047 (to M.P.C.). S.W.E. and R.T.L were both supported by Samuel and Althea Stroum Endowed Graduate Fellowships.

## AUTHOR CONTRIBUTIONS

J.S.G, R.T.L., M.P.C., D.A.G., and J.V. conceived the study. J.S.G., D.J.P., and A.G. performed mouse work, and J.S.G. prepared samples for metabolomics analysis. S.W.E. and R.T.L. processed tissue samples and collected and analyzed proteomics data. S.W.E. performed integrative analysis of all proteomics and metabolomics data. J.S.G. and S.M.J. performed the UCP1 mutant experiments in cell culture. M.M.P. contributed to data analysis. J.V. supervised the proteomics and global data analysis work. D.A.G. supervised mouse and validation work from J.S.G. and S.M.J. M.P.C. supervised work from D.J.P. and A.G. S.W.E., J.V. and D.A.G. wrote the paper, and all authors edited it.

## METHODS

### Mice

For all experiments, male C57BL/6J mice were used. Mice were housed in the Animal Medicine facilities of the UMMS in a clean room set at 22°C and 45% humidity under daily 12h light/dark cycles in ventilated racks with cages changed every two weeks. Mice were fed the facility chow diet. For the four weeks temperature acclimation experiments, mice at 10 weeks old were simultaneously housed in two rodent incubators (RIT33SD, Powers Scientific) within the Animal Medicine facilities of the UMMS. One of them had the temperature adjusted to 30°C (thermoneutrality group). Another incubator had its temperature decreased by four degrees weekly until reaching 6°C at which temperature the mice stayed for a week (severe cold group). Room temperature group mice were co-housed in the same facility as the mice in rodent incubators. Mouse cages were changed weekly using components pre-adjusted to temperature. No cage enrichment was used in this set of experiments. Mice were sacrificed at 14 weeks old in the morning in random fed conditions. For the “2-temp” experiment, mice were housed two per cage, with one nestlet per cage, at ambient room temperature. They were fed D12450J (Research Diets). At 8 weeks of age, mice in the thermoneutral group were transferred to a temperature and humidity controlled 30°C room. Cages were changed weekly using components pre-adjusted to temperature. Mice in the thermoneutral and room temperature groups were sacrificed at 12 weeks of age. All animal experiments were approved by the University of Massachusetts Medical School Institutional Animal Care and Use Committee.

### Analysis of Proteome and Acetyl-Proteome

For proteomics analysis, mouse interscapular BAT was homogenized in ice-cold lysis buffer using a plastic pestle in a microcentrifuge tube. The lysis buffer contained 8 M urea, 50 mM Tris pH 8.2, Roche complete mini EDTA-free protease inhibitor cocktail, phosphatase inhibitors (50 mM β-glycerophosphate, 1mM sodium orthovanadate, 10 mM sodium pyrophosphate, 50 mM sodium fluoride), and nicotinamide (class-III histone deacetylase inhibitor). Ground tissue was then sonicated for three cycles of 20 seconds each, chilled on ice for ∼10 min, and clarified by centrifugation at 16,000 x g at 4°C. Protein concentration was measured at this stage using the BCA Assay (Pierce). Protein cysteine residues were then reduced using 5 mM dithiothreitol (DTT) for 45 min at 55°C and alkylated using 15 mM iodoacetamide for 30 min at room temperature in the dark. Excess iodoacetamide was quenched with an additional 5 mM DTT at room temperature for 15 min. Samples were then diluted 5-fold in 50 mM pH 8.2 Tris, and digested with 5 μg/mL trypsin for 16 hours at 37°C, after which they were quenched by adding 10% trifluoroacetic acid to a pH of <2. Due to precipitated debris at the low pH, peptide samples were clarified by centrifugation at 3,500 x g at 4°C. Samples were then de-salted using Waters Sep-Pak tC18 cartridges (50 mg packing material), dried by vacuum centrifugation, and stored at −20°C until further use. For acetylation analysis, 0.5 mg of dried peptides were dissolved in 0.5 mL of ice-cold immuno-affinity purification (IAP) buffer (50 mM MOPS-NaOH, pH 7.2, 10 mM Na_2_HPO_4_, and 50 mM NaCl). Anti-acetyl-lysine agarose beads (ImmuneChem, lot #061214) were prepared by washing with cold IAP buffer. Approximately 25 μL of bead slurry was used per sample. Peptides and anti-acetyl-lysine beads were incubated on a rotating mixer for 2 hours at 4°C, washed five times with 1 mL ice-cold IAP buffer, followed by one wash with 1 mL phosphate buffered saline (Sigma-Aldrich). Acetylated peptides were eluted two times with 0.15% trifluoroacetic acid for a total of 100 μL, and desalted using four cutout layers of C18 Empore disks in a Stage-tip configuration^33^.

All liquid chromatography-tandem mass spectrometry (LC-MS/MS) experiments were done on a EASY-nLC 1000 (Thermo Scientific) coupled to a Q-Exactive mass spectrometer (Thermo Scientific). LC was performed with a 40 cm long, 100 μm inner diameter column packed in-house with Reprosil C181.9 μm beads (Dr. Maisch GmbH), with a column oven set to 50°C. Whole proteome and acetylation analyses were done using 90 minute LC-MS/MS runs, including column washing and equilibration. Prior to LC-MS/MS analysis, peptide samples were dissolved in a solution with 4% formic acid and 3% acetonitrile. For whole proteome analysis, samples were separated with a gradient of 9% to 32% acetonitrile in 0.15% formic acid over 62 minutes. For acetyl-proteome analysis, the gradient was 12% - 40% acetonitrile over 64 minutes. MS acquisition was performed using data-dependent acquisition with a “Top-20” method and 40 s dynamic exclusion. Precursor fragmentation was performed using higher-energy collisional dissociation with a 27% normalized collision energy. Whole proteome MS/MS acquisition was performed as follows: full-scan MS was acquired between 300-1500 m/z at 70,000 resolution, with a 3 × 10^6^ automatic gain control (AGC) target and 100 ms maximum injection time. MS/MS acquisition was performed with a 2 m/z isolation window at 17,500 resolution, with a 5 × 10^4^ AGC target and 50 ms maximum injection time. All MS data was collected in centroid mode. MS for acetyl-proteome analysis was performed similarly to the whole proteome analysis with the following differences: data-dependent acquisition was done with a “Top-10” method with 30 s dynamic exclusion, and the MS/MS maximum injection time was 100 ms. Proteome and acetylproteome MS analysis for the two-temperature validation experiment was done in technical duplicate. All MS data acquisition for proteomics was controlled using the XCalibur software version 2.2 (Thermo).

Raw files were converted to mzXML format and searched using Comet^34^ (version 2015025) against the mouse SwissProt database including isoforms (downloaded May 10, 2015). Methionine oxidation was treated as a variable modification, and carbamidomethylation of cysteine residues was treated as a constant modification. In the case of acetyl-proteome analysis, acetylated lysine residues were treated as a variable modification. Trypsin digestion was defined with cut-sites at lysine/arginine except when followed by proline. Precursor mass tolerance was set to 50 ppm, and fragment mass tolerance was 0.02 Daltons. Search results were filtered using Percolator^35^ (version 3.1.2) to obtain a list of peptide-spectrum matches at a 1% false discovery rate. Acetylation site localization was performed using an inhouse implementation of Ascore^36^ that admits any post-translational modification setting, where sites with an Ascore ≥ 13 (p ≤ 0.05) were considered localized. Quantification was performed using ThunderQuant, an in-house developed software for peptide quantification that relies on MS1 peak area integration. Peptide quantifications were pooled into protein quantifications, where the median peptide for each protein was taken to represent the protein. Protein quantifications were normalized to the total signal from each run. Acetylated peptides were pooled into acetylation isoforms, referred to “acetylsites” in the text for simplicity, if they have overlapping sequences and the same acetylation sites. If multiple unique peptides corresponded to an acetylation isoform, the median peptide was used for quantification. Acetylation isoforms were not normalized to the total signal from each run, because the total amount of acetylation, and the number of identified isoforms was seen to increase depending on the experimental condition. Differences between conditions were statistically determined by calculating two-tailed T-tests with unequal variance for each protein and acetylation isoform, and correcting the P-values with the Benjamini-Hochberg method. Proteins and acetylsites were labeled as significantly regulated if they are different between conditions at q ≤ 0.05, i.e. a 5% false-detection rate (FDR). Raw mass spectrometry data and search results are available on MassIVE (ID: MSV000082951).

### Metabolite profiling

Polar metabolites were analyzed at the Whitehead Institute Metabolite Profiling Core Facility. BAT samples were homogenized in four volumes of water using a TissueLyser II (Qiagen), and polar metabolites were measured using liquid chromatography coupled to tandem mass spectrometry (LC-MS/MS). A multiple reaction monitoring method was employed, where MS transitions, declustering potentials and collision energies were determined using reference standards. Metabolites were analyzed in either negative or positive ionization mode as previously described^37^.

Negative ionization mode data were acquired as follows. A 30μL aliquot of each homogenate was extracted using 120 μL of 80% methanol containing 0.05 ng/μL ^15^N_4_-inosine-, 0.05 ng/μL ^2^H_4_-thymine, and 0.1 ng/μL ^2^H_4_-glycocholate as internal standards (Cambridge Isotope Laboratories). The samples were centrifuged (10 min, 9,000 x g, 4°C) and the supernatants (10 μL) were injected directly onto a 150 x 2.0 mm Luna NH2 column (Phenomenex). Separation was performed with an ACQUITY UPLC system (Waters) running a hydrophilic interaction chromatography (HILIC) method. Initial mobile phase conditions were 10% mobile phase A (20 mM ammonium acetate and 20 mM ammonium hydroxide in water and 90% mobile phase B (10 mM ammonium hydroxide in 75:25 v/v acetonitrile/methanol) and the column was eluted at 400 μL/min using a 10 min linear gradient to 100% mobile phase A. Negative-mode MS/MS data were acquired using a 5500 QTRAP triple quadrupole mass spectrometer (AB Sciex). The ion spray voltage was -4.5 kV and the source temperature was 500°C.

Positive ionization mode data were acquired as follows. A 10-μL aliquot of each homogenate was extracted using nine volumes of 74.9/24.9/0.2 (v/v/v) acetonitrile/methanol/formic acid containing stable isotope-labeled internal standards: 0.2 ng/μL ^2^H_8_-L-valine-d8 (Isotec); and 0.2 ng/μL ^2^H_8_-L-phenylalanine (Cambridge Isotope Laboratories). Samples were centrifuged (10 min, 9,000 x g, 4°C) and supernatants (10 μL) were injected onto a 150 x 2.1 mm Atlantis HILIC column (Waters). Separation was performed with an 1100 Series pump (Agilent) and an HTS PAL autosampler (Leap Technologies). The column was eluted isocratically at a flow rate of 250 μL/min with 5% mobile phase A (10 mM ammonium formate and 0.1% formic acid in water) for 1 min followed by a linear gradient to 40% mobile phase B (acetonitrile with 0.1% formic acid) over 10 min. Positive ionization mode MS/MS data were acquired using a 4000 QTRAP triple quadrupole mass spectrometer (AB Sciex). The ion spray voltage was 4.5 kV and the source temperature was 450°C.

MultiQuant 1.2 software (AB Sciex) was used for automated peak integration and metabolite peaks were manually reviewed for quality of integration and compared against a known standard to confirm identity. Peak quantifications for each metabolite were normalized across all runs using the mass of the tissue sample. Differences between conditions were statistically determined by calculating twotailed T-tests with unequal variance for each metabolite, and correcting the P-values with the Benjamini-Hochberg method. Metabolites were labeled as significantly regulated if they are different between conditions at q ≤ 0.05, i.e. a 5% false-detection rate (FDR).

### UCP1 mutant plasmid construction and retroviral infection

Mouse UCP1 cDNA was subcloned into the pMSCV-puro-FLAG (C-terminal tagged) (Clontech) vector. UCP1 acetylation deficient-(KR) or mimetic-(KQ) mutant constructs were generated by site-directed mutagenesis using the NEBaseChanger tool for point mutation primers design. 293T cells (ATCC) were transfected with pMSCV-puro retroviral vectors expressing UCP1-Flag WT, UCP1-Flag 3KR, UCP1-Flag 3KQ, UCP1-Flag 4KR, UCP1-Flag 4KQ in combination with the retroviral packaging plasmids pCMV-VSVG (Addgene) and pCMV-Gag-Pol (Addgene). The culture media supernatant were added into target cells (293T) and followed by 24h incubation with Polybrene (8 ug/mL, Sigma). After incubation, the medium was replaced with complete medium. After 48h post infection, cells were trypsinized and subjected to puromycin selection.

### Gene expression and western blot analysis

For analyzing mRNA levels, total RNA was isolated using Qiazol (Qiagen) and RNeasy kit (Qiagen), cDNA synthesis was performed using High Capacity cDNA reverse transcription kit (#4368813, Applied Biosystems) and analyzed in a StepOnePlus real-time PCR machine. Primers for mouse UCP1 were 5′-CTGCCAGGACAGTACCCAAG and 3′-TCAGCTGTTCAAAGCACACA, and primers for TBP (a normalization control) were 5′-GAAGCTGCGGTACAATTCCAG and 3′-CCCCTTGTACCCTTCACCAAT. For western blot analysis, cells were harvested in cold PBS and lysed in lysis buffer (1% Triton X-100, 50mM Hepes at pH7.4, 150mM NaCl, 10mM beta-glycerol phosphate, 2mM EDTA, protease inhibitor cocktail). After elution by 5x sample buffer, the protein lysates were boiled followed by run (typically 10-20 ug per lane) in SDS acrylamide/bis-acrylamide gels, transferred to PVDF membranes and detected with indicated antibodies. The UCP1 antibody (ab10983) is from abcam. The Flag antibody (CAT# F1804, clone M2) is from Sigma. The α-tubulin antibody (CAT# 2125, clone 11H10) is from Cell Signaling technologies. All antibodies have validation statements on their manufacturers’ websites.

